# Time-series modeling with neural flow maps

**DOI:** 10.1101/2025.06.18.660388

**Authors:** Bingxian Xu, Zoey E. Ho, Yitong Huang

## Abstract

Constructing mathematical models from data is fundamental for understanding complex systems across scientific disciplines. However, real-world data often pose challenges such as irregular sampling, sparsity, and noise, that hinder the development of accurate, mechanistic models. In this work, we present a deep learning framework that directly reconstruct flow maps from data, assuming only that the observed patterns arise from an autonomous dynamical system. We demonstrate that our method accurately captures system dynamics across diverse settings, even with limited and irregularly sampled training data. When applied to the circadian transcriptomic time series data, it generates biologically valid predictions by integrating information across multiple organs. By parameterizing the full dynamical system, our proposed approach enables efficient computation of time derivatives and Jacobians directly from data, offering a powerful tool for analyzing and interpreting high-dimensional biological systems.

## 1 Introduction

Data-driven modeling for complex systems holds great promise for accelerating scientific discoveries, ranging from uncovering new equations for atmospheric dynamics [1, 2] to predicting tumor growth [3, 4, 5]. Such models can aid in identifying key components of a system and informing the design of control strategies. However, constructing accurate and interpretable models becomes increasingly challenging when data are high-dimensional and sparse.

Initial efforts in this field used symbolic regression methods to discover the governing equations [6, 7]. While powerful in principle, these methods face challenges in practice due to their high computational costs. More recently, approaches such as the sparse identification of nonlinear dynamics (SINDy) [8] have gained popularity because of their much improved computational efficiency, achieved through the introduction of a predefined library of candidate functions. These strategies have also been applied to biological systems [9], where the governing equations are traditionally hand-curated based on prior knowledge and then fitted to experimental data [10, 11, 12, 13]. However, these methods face substantial limitations in high-dimensional settings. SINDy-like approaches require accurate estimation of time derivatives, which is often impractical when data are sparse or noisy. Moreover, the sheer number of variables in such systems often precludes the use of hand-curated models or the construction of a suitable candidate function library.

These challenges motivate the development and application of modeling approaches that are agnostic to the detailed mechanisms of the system, methods that require neither prior knowledge nor a pre-defined set of governing equations. For example, both PHOENIX [14] and RNAForecaster [15] can be trained with sparse high-dimensional transcriptomic time series without prior knowledge by parametrizing the underlying dynamical system with neural networks. While using neural differential equations (neural ODEs) [16] allowed one to fit complex dynamical systems, it has been shown that they often get trapped into local minima and can fail even on relatively simple oscillatory patterns [17]. Moreover, their training requires costly time-integration during both the forward and backward passes, making them less practical for systems with multiple time-scales. These limitations, inherent to vector field based parametrization, can be avoided by instead learning a flow map, which can theoretically map the system forward arbitrarily far in time with one single forward pass. While this may be not always achieved in practice, flow maps still offer much faster prediction by directly approximating the integral form of the underlying differential equations [18].

Flow map learning have been extensively explored and applied in various contexts [19, 20, 21, 22, 23, 24, 25], including biological [24], chaotic [25] and stochastic systems [21]. However, to our best knowledge, most existing parametrizations of flow maps are restricted to predicting the system’s state at a fixed time interval [19, 20, 21, 22, 23, 24, 25], with intermediate states typically inferred via interpolation. In other words, their reconstructed flow maps are approximations of the “true” dynamics only for a single predefined time step Δ*t*. As a result, such models cannot directly compute key quantities of the underlying dynamical systems, such as the vector field or the Jacobian matrix.

To bridge the gap between data-driven flow maps with mathematical theory, we build on existing work [18, 26] and propose novel strategies to enforce the mathematical properties of flow maps and utilize their advantages for modeling dynamical systems. With time encoding, our framework can predict the system’s future state via a single forward pass without performance degradation at large time intervals. In addition, we apply a multi-scale regularization to enforce the semi-group property, a key characteristic of flow maps. These design choices enable us to recover the underlying vector field directly from the reconstructed flow map. We show that our proposed architecture can reliably capture key features of the underlying system even when trained on sparse time series data. When only cross-sectional data are available, our approach can efficiently learn the time evolution of probability distributions. Finally, we apply our method to sparse and noisy transcriptomic time series, showing that it can reconstruct gene expression dynamics and identify potential regulatory genes through analysis of the Jacobian matrix.

## 2 Methods

### 2.1 Setup

Given a system of ordinary differential equations (ODEs) of the form

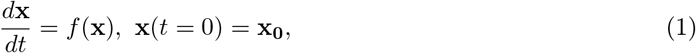

where **x** ∈ ℝ^*n*^ is the set of state variables of the system and *f*: ℝ^*n*^ *→*ℝ^*n*^ is a function that governs their rate of change. The general solution to Equation 1 can be expressed as

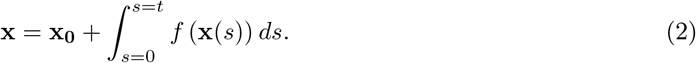

The flow map operator of this system is a function Φ that describes the evolution of state variables over time. Specifically, it maps the initial condition **x**_**0**_ to the state **x**(*t*) at time *t*:

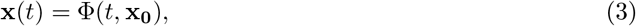

which is denoted as Φ_*t*_(**x**_**0**_) for simplicity. The flow map operator must satisfy two important properties:

1. Φ_0_(**x**) = **x** (Identity mapping)
2. 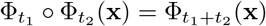 (Semigroup)

### 2.2 Flow architecture with time embedding

The resemblance between Equation 2 and a single residual layer motivates the use of the residual network (ResNet) [27] to parameterize the flow map operator [19, 20, 21, 22, 23, 24, 25]. Specifically, the basic structure is defined as follows [18, 26]:

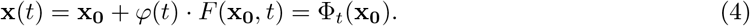

Here, we choose *φ*(*t*) = tanh(*t*) to enforce identity mapping, and parameterized *F* (**x**, *t*) with neural network(s). The functional form of *φ*(*t*) is not necessarily unique; for instance, setting *φ*(*t*) = *t* has been proposed [18]. However, directly inputting *t* into the neural network *F* (*·*,*·*) may impair the model’s ability to interpret time meaningfully, especially when *t* is too small or large. For example, when working with datasets with multiple time scales, Chen et al. [18] used log_10_(*t*) instead of *t* to address this issue.

In our setup, we leverage the Bochner’s Theorem and project time into a bounded, higher dimensional space. Let *κ* be some kernel such that *κ*(*t*_1_, *t*_2_) measures the distance between the projection of *t*_1_ and *t*_2_ in the high-dimensional space. Intuitively, it is desirable for this kernel to be translation invariant, i.e., for *κ*(*t*_1_, *t*_2_) to depend only on the difference *t*_1_*− t*_2_, rather than the specific values of *t*_1_ and *t*_2_. The Bochner’s theorem, stated below, provides key insights into the construction of such time embeddings.

#### THEOREM 1

(Bochner’s Theorem). *A continuous, translation-invariant kernel κ*(*x, y*) = *g*(*x − y*) *on* ℝ^*d*^ *is positive definite if and only if there exists a non-negative measure on* ℝ *such that g is the Fourier transform of the measure*.

Bochner’s Theorem states that *κ* can be expressed in the form:

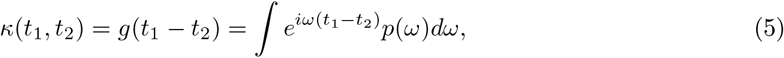

where *p*(*ω*) is the probability measure on ℝ. Taking only the real part of equation 5, we obtain

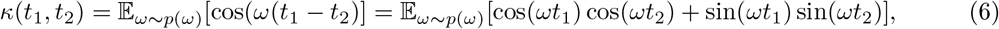

which implies a feature map of the form [28],

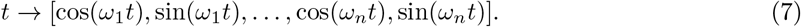

In our setup, we learn the set {*ω*_*i*_}directly during the optimization process. The encoded time coordinates are then concatenated with the state variables and passed as input to the neural network.

### 2.3 Semigroup-enforced learning

One of the key advantages of constructing a flow map is its independence from numerical integration. In principle, this allows the model to predict the system state **x** at any future time *t* in a single forward pass:

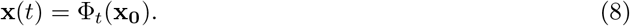

However, leveraging this one-shot prediction capability requires the model to be trained with appropriate regularization to ensure stability and generalization across time scales—a direction that remains relatively underexplored in the existing literature.

As demonstrated in prior work, flow map learning frameworks are typically implemented using a recursive ResNet, where the model structure mirrors the forward Euler method with a fixed time step. In this formulation, the learned dynamics effectively approximate a system of difference equations that propagate forward in time:

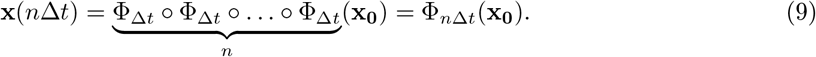

However, reliance on this forward scheme constrains the training data to be sampled at fixed time intervals, which can be difficult to achieve in real world settings, especially when large volumes of data are required. To maximize the flexibility of the reconstructed flow map, it is desirable for the model to adapt to a range of time steps Δ*t*, thereby underscoring the importance of enforcing the semigroup property of the flow map operator.

Recently, it has been proposed that the semigroup property can be enforced by minimizing a two-step loss function [18, 26]:

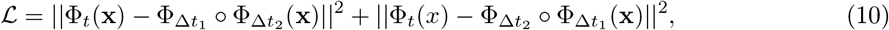

where *t* = Δ*t*_1_ + Δ*t*_2_. However, we empirically observed that this formulation becomes less effective when the training data is sparsely sampled. To address this limitation, we propose a multi-step approach to enhance the semigroup property by minimizing:

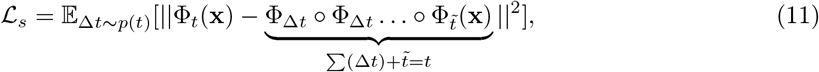

where 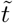 denotes the difference between *t* and ΣΔ*t*. During training, we sample Δ*t* from a uniform distribution, enabling the model to learn across different time scales.

In addition, we incorporate a data-driven reconstruction loss to ensure consistency with the observed data. Given training data **X**_**i**_ ∈ ℝ^*v×n*^ with corresponding time coordinates *t*_*i*_, where *v* is the number of features and *n* the number of state variables, we define the reconstruction loss as:

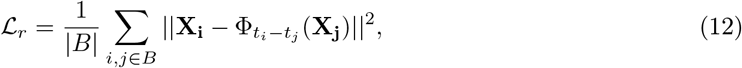

where *B* is a randomly sampled batch from the training data such that such that *t*_*i*_ *> t*_*j*_. The total loss used to train our network combines the semi-group loss and the reconstruction loss, ℒ= ℒ_*s*_ + ℒ_*r*_.

## 3 Results

We demonstrate the algorithm on a number of example systems to highlight its diverse applications, ranging from time-series reconstruction from canonical dynamical systems, to optimal transport problems, and finally to experimental data from circadian clocks.

### 3.1 Time series reconstruction

#### 3.1.1 Van der Pol oscillator

The first example illustrates the performance of our algorithm using synthetic data generated from the Van der Pol oscillator. Because biological data collected in experiments are often limited and sparse, our goal is to show that the algorithm can accurately reconstruct time series from a single trajectory **x**(*t*), even when observations are sparse and of limited duration. In this context, successful learning is defined as the algorithm’s ability to capture the general structure of the stable manifold, rather than its performance across the entire phase space.

In the example shown in Figure 1A, data are collected from *t* = 0 to *t* = 20 with a time step of Δ*t* = 0.5 for one initial condition. Figure 1B shows that our proposed algorithm successfully reconstructs the underlying vector field by computing 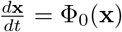. Furthermore, reconstructing time series iteratively using Equation 9 with varying values of Δ*t* indicates that the semigroup property of the flow map operator is preserved (Figure 1C). Compared to previously proposed flow map learning algorithms that use a two-step loss function [26, 18], our implementation of multi-step loss regularization yields significant improvement in preserving the semigroup property under sparse sampling conditions, that is, as the total time interval *t*_1_ + *t*_2_ increases (Figure 1D, S1A).

**Figure 1:**
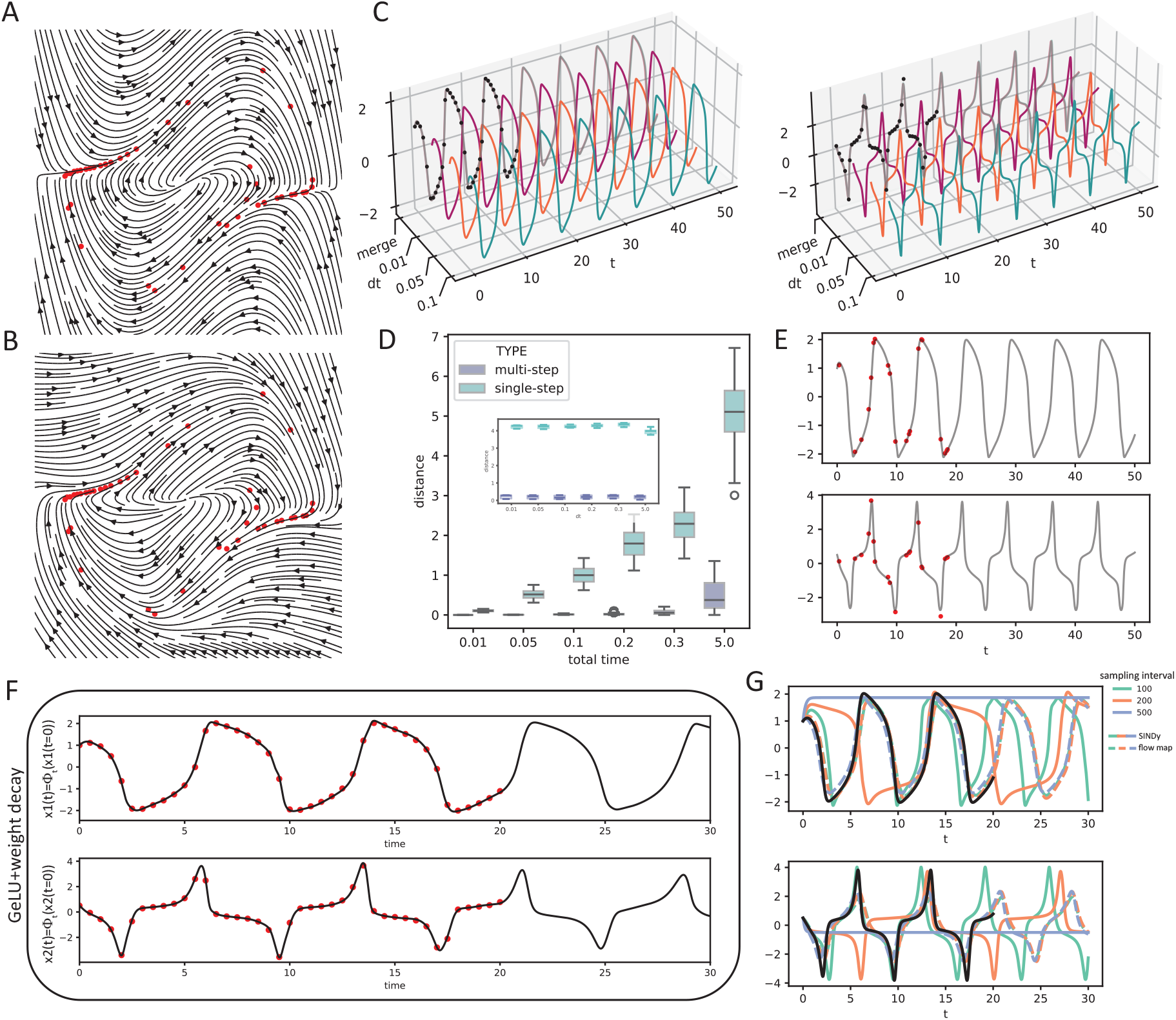
Flow map learning for the Van der Pol oscillator. A: Phase portrait of the true Van der Pol oscillator, with training data shown as red dots. B: Phase portrait of the learned system with training data (red dots) overlaid. C: Reconstructed time series using the iterative method for different time intervals (training data shown in black). D: Evaluation of the semigroup propety for models trained with different semigroup loss, quantified by the distance between 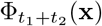 and 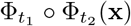, where *t*_1_ + *t*_2_ denotes the total time. Inset: Distance between Φ_*dt*_ ○ Φ_*dt*_ … ○ Φ_*dt*_(**x**) with a fixed total time of 10. E: Reconstructed time series using the iterative method with irregularly sampled training data (red). F: Reconstructed time series obtained from a single forward pass. G: Reconstructed trajectories (colored) and true solution curves (black) under various sampling intervals and a noise level of 0.5.

To fully leverage the one-shot prediction capability of the flow map operator, i.e., constructing the flow map in a single forward pass, we implemented several regularization techniques. A key observation is that combining the GeLU activation function with weight decay outperforms using GeLU, weight decay, or Lipschitz constraints individually (Figure 1F, S1B), confirming that appropriate regularization is necessary for ensuring the model’s generalization across time scales.

The proposed flow map learning algorithm is also competent under irregular sampling, a scenario not previously addressed by existing methods. As shown in Figure 1E, the algorithm accurately recovers the underlying dynamics using only 20 irregularly sampled observations collected between *t* = 0 and *t* = 20.

We then investigate the effect of sampling frequency and observational noise on model performance. Specifically, we introduced additive noise drawn from 𝒩 (0.5, 1) (a standard normal with with mean 0.5 and unit variance) to the observations. Figure S2 shows that an existing method using sparse regression for recovering dynamical systems, SINDy, is highly sensitive to both decreased sampling frequency and increased noise. While SINDy can accurately reconstruct the oscillatory dynamics when sampling frequency is high, it fails to capture the oscillation at all when observations are sparse and noisy. In contrast, our proposed algorithm consistently recovers the underlying oscillatory dynamics with good accuracy, regardless of the sampling frequency (Figure 1G and S2).

#### 3.1.2 Chaotic oscillators

Next, we evaluate the performance of our method for chaotic systems. As an illustrating example, we generate a single solution trajectory from the Lorenz chaotic attractor for one initial condition. Specifically, two datasets with different sampling frequencies were constructed: a short, high-frequency dataset with timestep *dt* = 0.005 over the interval *t* = 0 to *t* = 10 (Figure 2A), and a longer, low-frequency dataset with timestep *dt* = 0.1 over the interval *t* = 0 to *t* = 30 (Figure 2B). It is worth pointing out that the both datasets used for training are substantially smaller than those typically used in the literature for chaotic systems (e.g., *dt* = 0.01 over *t* = [0, 10, 000] in [25], and *dt* = 0.001 over *t* = [0, 100] in [8]).

**Figure 2:**
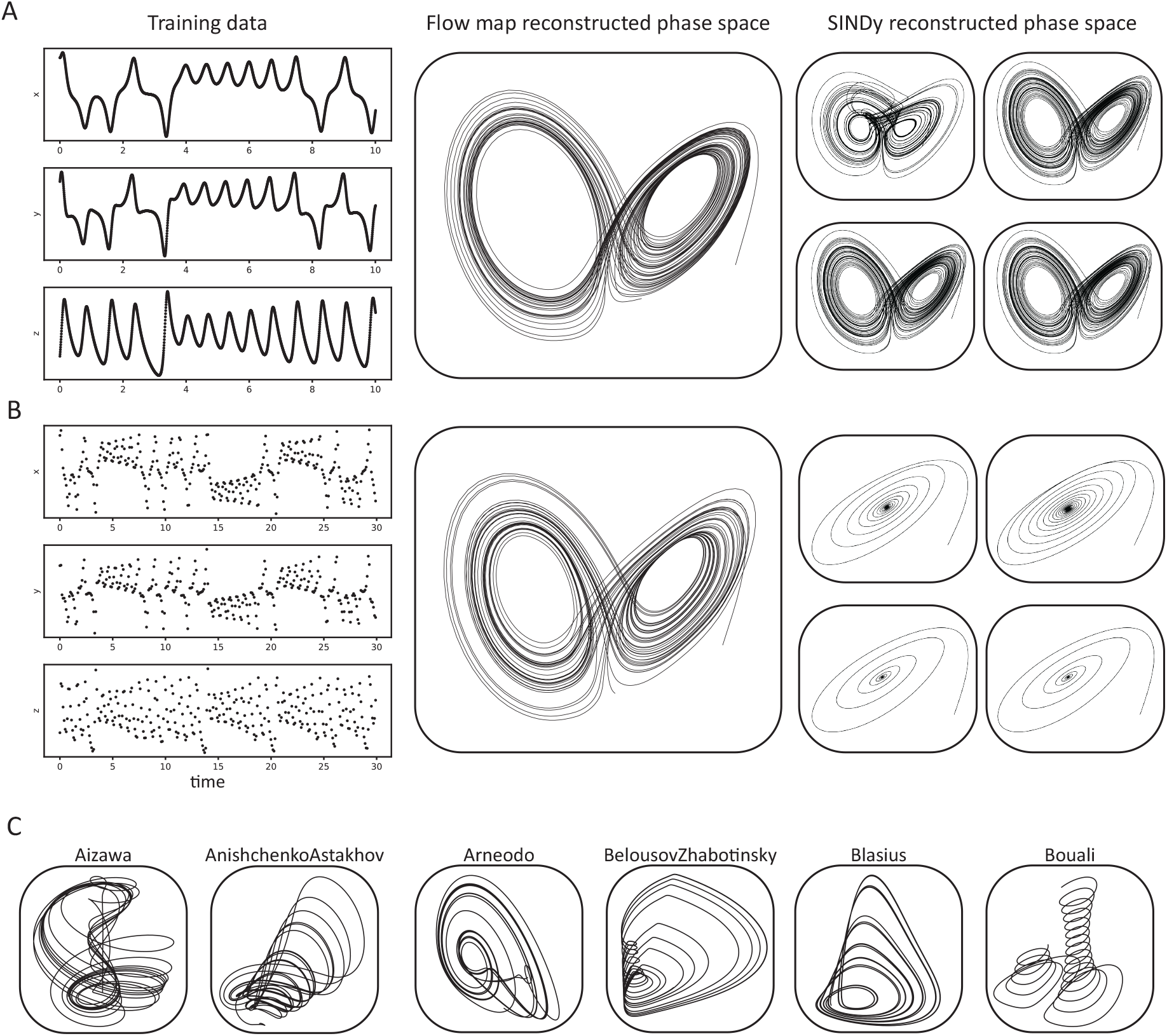
Flow map learning for Chaotic oscillators. A: Comparison of high-resolution training data (first column), flow map generated phase space (second column), and SINDy generated phase space (third column). B: Comparison of low-resolution training data (first column), flow map generated phase space (second column), and SINDy generated phase space (third column). For the SINDy generated phase space in the third column of panel A and B, each set contains four subpanels, showing results obtained using polynomial libraries of increasing power: degree 5 (top left), degree 4 (top right), degree 3 (bottom left), and degree 2 (bottom right). C: Examples of chaotic oscillator trajectories reconstructed using the flow map learning framework.

The performance of SINDy is known to depend heavily on the choice of coordinates and the function library. While SINDy with a polynomial basis up to degree 4 can accurately recover the Lorenz attractor when trained on the high-frequency dataset, it fails to capture the attractor shape when trained on the sparse, low-frequency dataset, regardless of the basis functions used. This observation, along with our earlier results for the Van der Pol oscillator in Figure 1G, supports findings in the literature that highlight SINDy’s sensitivty to data quality and sampling density [29, 30, 31, 32]. In contrast, our algorithm demonstrates robustness in model recovery. Figure 2 shows that it successfully captures the shape of the Lorenz attractor in both high- and low-frequency scenarios, despite the limited data. Additionally, Figure 2C shows that our algorithm is generalizable for recovering other three-dimensional chaotic systems. Details of data generation for these chaotic systems are provided in the supplement.

### 3.2 Optimal transport

We next consider a setting in which our algorithm reconstructs dynamics that are inaccessible to existing system identification methods. Most existing approaches focus on longitudinal time series; however, many biological data, such as single-cell RNA sequencing, capture high-dimensional measurements from individual cells at a single time point. While these cross-sectional measurements provide rich information, they lack temporal continuity and are challenging for traditional methods.

To address this issue, we modify our flow map learning algorithm to solve a distribution-matching problem, which can be formulated as an optimal transport task:

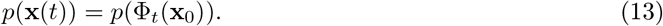

Here, *p*(*·*) denotes a probability distribution, **x**(*t*) denotes the states of samples collected at time *t*, and **x**_0_ denotes the states measured at time 0. In this setting, the number of samples collected at different times need not be equation, and the temporal trajectories of individuals are unknown. As a result, standard loss functions, such as mean squared error, are not application. Instead, we construct flow maps by minimizing the distance between distributions, measured by the sliced Wasserstein’s distance:

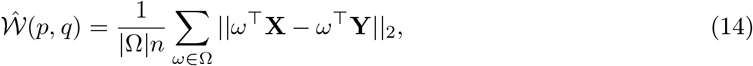

where *p* and *q* denotes probability distributions from which **X** and **Y** are sampled (assume *ω*^T^**X** and *ω*^T^**Y** are sorted in ascending order), *n* is the dimension of the system and Ω is a set of random projection directions. We then revise the reconstruction loss ℒ_*r*_ and semigroup loss ℒ_*s*_ as:

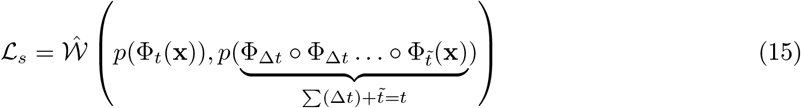

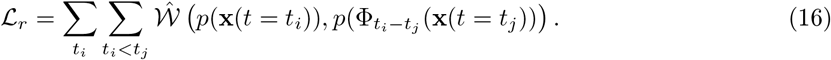

To demonstrate our approach can learn the evolution of distributions, we start with simulating training data in which a central “source” distribution branches into multiple “target” distributions. Figure 3A shows that our algorithm was able to successfully learn the branching dynamics in this setting. Notably, we find that distribution matching typically significantly fewer training iterations compared to learning from longitudinal time series data, likely due to the reduced complexity of motion in phase space.

**Figure 3:**
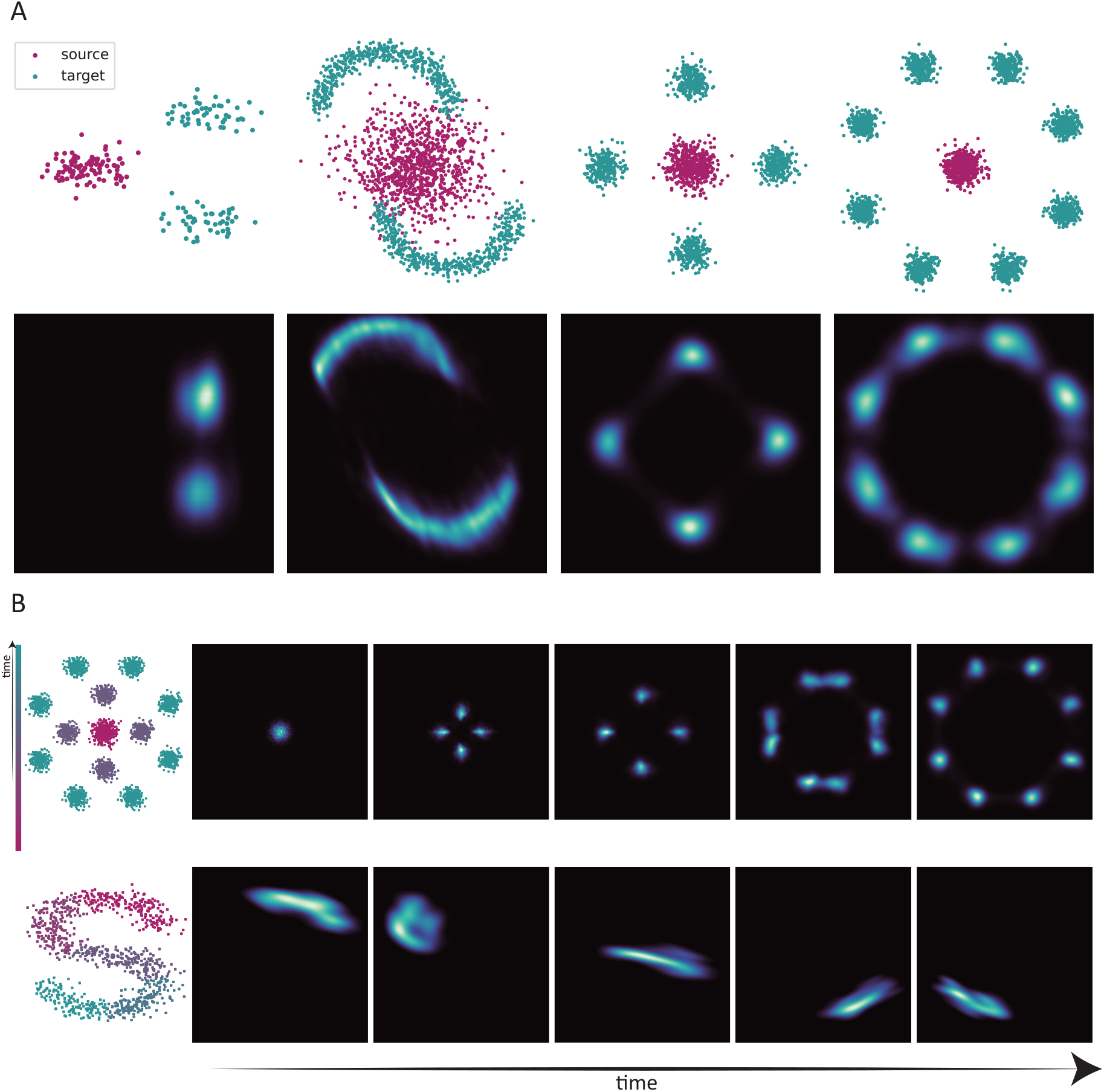
Optimal transport with flow map. A: Training data (top row) and reconstructed distributions (bottom row) with increasing complexity. B: Time evolution of probability distributions predicted by the flow map model.

We then evaluate the method on more complex cross-sectional data involving more than two time points and again observe good reconstruction performance (Figure 3B). Importantly, the flow maps learned in our approach correspond to an underlying autonomous dynamical systems. This contrasts with other methods, such as the continuous normalizing flow [16] or trajectory net [33], which rely on explicitly time-dependent flows. Finally, by using the sliced Wasserstein’s distance, our approach remains effective even when the source and target distributions cannot be easily parameterized, offering flexibility for application to real-world biological data.

### 3.3 Real-world application in circadian rhythms

The results have so far shown that our algorithm can recover dynamics from synthetic data. We next apply it to a real-world example: time-series transcriptomic measurements collected from multiple mouse organs [34]. Circadian rhythms have long been a subject of interest in the biological literature; the ability of high-throughput gene expression profiling has opened new avenues for uncovering the molecular basis of circadian regulation and its implications for health. However, a major difficulty in this context is the experimental burden and high cost of sequencing, which limits the ability to collect samples around the clock. Furthermore, gene expression data are inherently noisy due to biological variability and technical noise introduced during sample processing. By applying our algorithm on the mouse transcriptomic dataset, we show that: (1) our method can recapitulate expression profiles for individual genes in each organ, and (2) the algorithm reveals a shared gene regulatory network structure that reflects biologically meaningful coordination across tissues.

To assess whether our framework can be applied to real biological systems, we acquired a multi-organ circadian transcriptomic time series dataset from [34]. This dataset exhibits several of the challenges discussed in previous sections— for example, it is sampled every two hours over a 48-hour period, making it both short in duration and limited in temporal resolution. We first apply our framework to the adrenal gland, selecting the 50 genes with the greatest circadian variation as the state variables, **x**. Using data from the first 24 hours as the training set, we find that our model fits the data well and is able to “generates” sustained oscillation, thereby capturing the underlying circadian dynamics of the system (Figure 4A).

**Figure 4:**
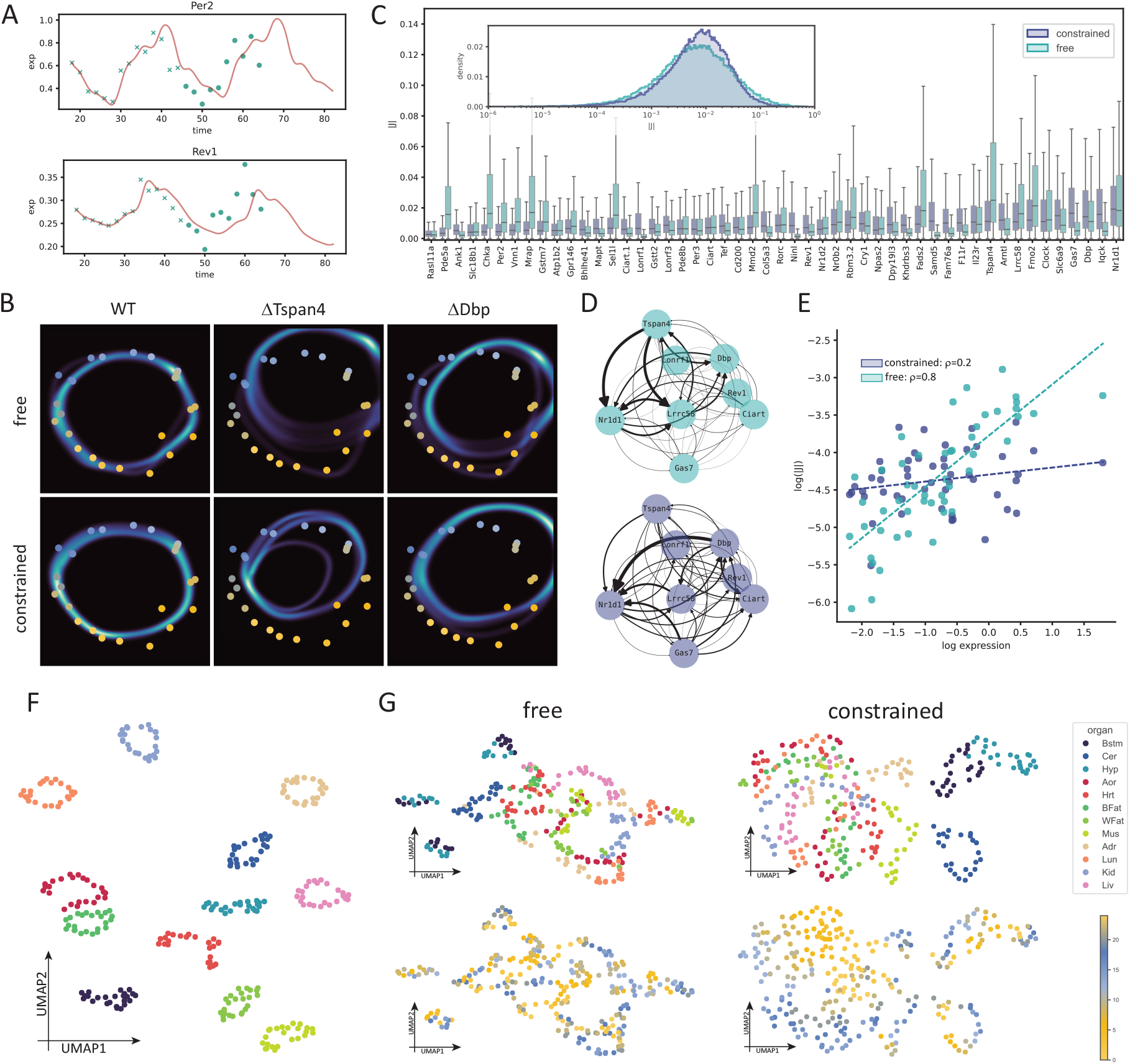
Flow map learning of transcriptomic time series. A: Example gene expression dynamics generated by the model trained only on the first circadian cycle (crosses). B: Observed gene expression data projected onto the first two principal components and colored by sampling phase (dots). The heatmap shows model-predicted dynamics projected into the same PCA space. WT: “wild type” system experiencing no gene knock out. ΔTspan4 and ΔDbp refers to systems where gene Tspan4 and Dbp is constantly set to 0. C: Boxplot of gene-wise regulatory power in the adrenal gland, quantified by the absolute values of the entries of the Jacobian matrix. (Inset: Distribution of regulatory powers of all genes for the constrained and the free model). D: Directed regulatory networks for Nr1d1 and Tspan4 in the adrenal gland, extracted from the Jacobian matrix of the constrained and free model. Edge weights are computed by average absolute Jacobian entries across all sampled states. E: Relationship between each gene’s total regulatory power (row-wise absolute sum of the Jacobian matrix) and its average expression level in the adrenal gland. F: UMAP embedding of gene expression profiles from all 12 sampled organs. G: UMAP embedding of gene regulatory rules, derived from the Jacobian matrix of the constrained and free models, colored by organ of origin (top) and sampling phase (bottom).

Next, we seek to exploit the fact that this dataset contains measurements from multiple organs, each occupying a distinct region in gene expression space. Using the same 50 genes selected from the adrenal gland, we tested whether incorporating dynamics from other organs could offer additional insight. To this end, we trained two models, one *constrained* by data from additional organ, and one *free* of them. Both models fit the data well (Figure 4B, WT column).

We then computed the Jacobian matrix from these models, *J*, where the entry, 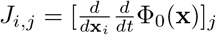, quantifies how changes in gene **x**_*i*_ affects the dynamics of gene **x**_*j*_. We used the absolute value of the sum of each row of the Jacobian matrix as a proxy for each gene’s regulatory power. Interestingly, in the constrained model, the gene with the highest regulatory power was Nr1d1, a well-known core circadian regulator (Figure 4C). In the free model, while Nr1d1 remained influential, Tspan4 emerged as the dominant regulator.

To further explore differences between these models, we constructed a small weighted directed network composed of the top targets of Nr1d1 and Tspan4 (Figure 4D). In the free model, Tspan4 exerted a strong influence on Nr1d1, an effect that was markedly reduced in the constrained model (Figure 4D). Similarly, simulation of a Tspan4 knockout had a larger impact on the system defined by the free model than in the constrained one (Figure 4B). Conversely, Dbp showed stronger influence on Nr1d1 in the constrained model, again confirmed by knockout simulation (Figure 4B, D).

Further analysis revealed a key distinction between the models: in the free model, the entries of the Jacobian matrix were strongly correlated with gene expression levels in the adrenal gland, suggesting that high expression levels were conflated with regulatory importance (Figure 4E, free model: Pearson correlation coefficient *ρ* = 0.8, *p* = 9 *×* 10^*−*12^, constrained model: *ρ* = 0.2, *p* = 0.15). In contrast, the constrained model’s Jacobian was not correlated with the overall gene expression (*ρ* = 0.13, *p* = 0.36). This suggests that, in the free model, training on a highly confined region of state space biases the learned dynamics, while the additional organ data in the contrained model acts as a form of regularization. This effect is further supported by the distribution of Jacobian entries (Figure 4C, inset), which shows that genes with high expression in the free model gained disproportionately high regulatory power, while this bias was attenuated in the constrained model. These observations indicate that training on a broader sampled state space, achieved here through multi-organ data, helps mitigate overfitting and prevents the model from equating expression magnitude with regulatory influence. This regularizing effect aligns with prior findings on improving robustness in dynamical system models [35, 36]. Indeed, although both models perform comparably in terms of data fit, the free model exhibits greater signs of overfitting (Figure S3).

To see whether our earlier conclusions hold when expanding the dimensionality of the state space, we trained an additional model using the top 500 circadian genes. The resulting model was able to successfully reconstruct the covariance structure of the data (Figure S4). Surprisingly, we found that the increasing the number of genes led to a further reduction in the absolute values of the entries in the Jacobian matrix (Figure S5). Despite this, the genes with the highest regulatory power included Nr1d1 [37], Fabp7 [38, 39], Dbp [40, 41] and Per2 [42], all of which are well-established genes in circadian regulation [43].

Together, these results demonstrate the viability of using biological data to constrain mathematical models of gene regulatory dynamics. Even when trained on a small subset of detected genes, incorporating information from multiple organs effectively regularizes the Jacobian matrix, and the resulting model identifies biologically meaningful circadian regulators as key components of the system. This supports the idea that the breadth of sampled conditions can improve model robustness and interpretability.

### 3.4 Unifying tissue specific gene regulatory networks

Gene regulatory networks (GRNs) are usually represented as directed or undirected networks, where nodes correspond to genes and edges denote gene-gene interactions. Despite a long history of development [44, 45, 46], data-driven reconstruction of GRNs remains an active area of research [47, 46]. To our knowledge, GRNs are typically considered to be highly context specific [48, 49, 50], often treated as “discrete” and independent entities. As a result, it is generally difficult to infer a GRN under one condition based on knowledge of another, and most GRNs are not designed to reconstruct or simulate underlying gene expression dynamics. In contrast, a flow map fully defines the behavior of a dynamical system and thus implicitly encodes the interactions each gene participates in. When the system resides in different regions of the gene expression space, such as those corresponding to different tissues, context-specific GRN may be represented by the Jacobian matrix of the flow map evaluated at those locations. For instance, if *J*_*ij*_ *>* 0, gene *i* is inferred to positively regulate gene *j*, and vice versa. By computing these Jacobians at various sampled states and vectorizing them, we can apply dimensionality reduction techniques such as UMAP [51] to visualize how regulatory interactions change across covariates like tissue type or time.

Using the top 50 circadian genes from the adrenal gland to define our state space, we found that expression profiles from different organs occupied clearly distinct regions within the gene expression space (Figure 4D). When applying the free model, we observed that the gene-gene interactions inferred from the flow map Jacobians were highly tissue specific (Figure 4G). However, in the constrained model, trained across all organs, the low-dimensional projections of flow map Jacobians derived from visceral organs formed a ring-like structure and were separate from the brain. Notably, when we colored the UMAP of gene-gene interactions by sampling time, we observed a clear temporal progression (Figure 4G). This findings suggest that, while gene expression profiles are highly tissue-specific (Figure 4F), the underlying mechanism, captured by the Jacobian of the flow map, can be shared across tissues and evolve coherently in time.

## 4 Discussion

In this work, we developed a deep learning framework to directly reconstruct flow maps from data. Our framework incorporates several key features that exploit the unique advantages of flow maps. First, unlike previous approaches [19, 20, 21, 22, 23, 24, 25], our method is capable of handling sparse and irregularly sampled data, a common challenge in real-world settings. Moreover, we introduced a novel loss function to strictly enforce the semigroup property, ensuring temporal consistency in learned dynamics. Our use of time encoding inspired by Bochner’s Theorem [28] further enables efficient “oneshot” time series generation described by equation 8.

A main advantage for parameterizing a dynamical system via a flow map, rather than a system of ODEs, is that it avoids the need for numerical integration, an operation more complex and error-prone than numerical differentiation. NeuralODEs are known to be difficult to train even for simple systems due to their tendencies to stay in local minima [17]. Yet flow maps and ODEs need not be seen as fundamentally distinct: after learning a flow map Φ_*t*_, one can estimate the corresponding vector field 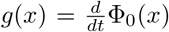, making our framework well-suited for hybrid approaches that rely on accurate derivative estimates, such as SINDy.

We illustrated the utility of our method across a diverse set of systems. With the Van der Pol oscillator, we showed that our framework performs well even with limited training data. In the Lorenz system, where time-series prediction is less relevant than capturing underlying structure, our approach better recovered the attractor dynamics than alternative methods (Figure S2). With the circadian transcriptomic data, our framework successfully reconstructed gene expression dynamics and revealed biologically meaningful structure in the Jacobian of the flow map’s time derivative. Notably, this structure only emerged when the model was trained with data from multiple organs. In particular, known circadian genes, such as Nr1d1 and Dbp, emerged as key drivers whose perturbation substantially altered the system’s behavior.

Visualizing gene-gene interactions derived from by the Jacobians on a lower-dimensional space revealed a time-dependent structure in the constrained model. This implies that system responses may vary depending on when a perturbation occurs, a result that aligns with findings from circadian medicine. [52, 53, 54]. In addition, the observation that Jacobians from different organs collapse onto a ring-like structure despite tissue-specific expression suggests that the underlying system is approximately linear 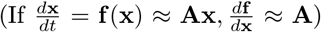. This hypothesis is supported by recent studies showing that phase difference of circadian genes increases as their distance along some gene regulatory network increases [55]. If the system is not largely linear, phases of interacting genes would be unrelated. Taken together, our result serves as a proof of principal that dynamical models become more biologically informative when constructed using data from distinct regions of the gene expression space. Our approach also generalizes to settings where only cross-sectional data is available. We showed that our approach can be used to construct transport maps between distributions, exemplifying its versatility. This suggests our framework could extend to single cell RNA-seq data in the future.

Despite these promising results, enforcing mathematical properties of flow maps remains challenging. While our semigroup loss performed well on simpler systems like the Van der Pol oscillator, it was less effective in chaotic systems like the Lorenz system. On the other hand, while a number of ways have been proposed to enforce the invertibility of neural networks [56, 26]. These methods are either inapplicable, or was observed to severely reduce the performance of the model. Another open problem is defining more suitable loss functions in the furture. For instance, when modeling gene expression data with a wide dynamic range, standard mean squared error can disproportionately emphasize highly expressed genes, underrepresenting the importance of lowly expressed but functionally critical ones. Nevertheless, in an era of increasingly large and complex datasets, when hand-writing equations becomes infeasible, our framework offers a flexible, interpretable alternative. By learning a flow map that governs system behavior, we can identify key components, predict responses to perturbations, and ultimately aid in the design of intervention strategies.

