## supplemental figures and notes for "Time-series modeling with neural flow maps"

### 5 Supplemental material

#### 5.1 Model architecture and training

By default, we used a series of three residual networks to parametrize the flow map. Each residual network contains a four-layer neural net with 100 neurons in each layer. When dealing with more complex time series such as the chaotic oscillator, we used 600 neurons per layer.

Optimization of the neural network parameters is done with ADAM with $\beta = [0.8, 0.9]$ and a weight decay of $10^{-5}$. During training, the learning rate is set to change between $10^{-5}$ to $5 \times 10^{-4}$ using a cyclic scheduler.

#### 5.2 Constructing training data

##### 5.2.1 Van der Pol oscillator

Tthe Van der Pol oscillator is defined as:

$$\begin{cases} \frac{dx}{dt} = y \\ \frac{dy}{dt} = 2(1 - x^2)y - x \end{cases}$$

After solving the equations with Euler's method with a time step of 0.001, the training data is sampled from the numerical solution with an sampling interval of 500. This means that for every 500 points on the numerical solution, we only take one into the training set. Therefore, decreasing sampling interval suggests enhanced temporal resolution.

When testing how our method works under the influence of noise. We inject standard white noise to the solution curve with a fixed standard deviation. In the main text, we refer to the standard deviation of the injected noise as noise level.

##### 5.2.2 Chaotic oscillators

The Lorenz system we used is defined below:

$$\begin{cases} \frac{dx}{dt} = 10(y - x) \\ \frac{dy}{dt} = x(28 - z) - y \\ \frac{dz}{dt} = xy - \frac{8}{3}z \end{cases}$$

It is solved with Euler's method with a time step of 0.0001. Solutions to the additional chaotic oscillators were generated using the `dysts` [57] package. The training data consists of 1000 time points of a single solution curve.

#### 5.3 Assessing the semigroup property

The semigroup property was assessed by two ways, both of which used 10000 randomly sampled points from the square bounded by $x \in (-2.5, 2.5)$ and $y \in (-4, 4)$. In our first approach, we compared the difference between $\Phi_t(x)$ and $\Phi_{t1} \circ \Phi_{t2}(x)$ where $t$ is fixed (Figure 1D). In the second approach, we compared the difference between $\Phi_t(x)$ and $\Phi_{dt} \circ \Phi_{dt} \ldots \Phi_{dt}(x)$ where $dt$ is fixed and $t$ is set to 10.

#### 5.4 Assessing the goodness of fit

The goodness of fit for the reconstructed Van der Pol oscillator is quantified by its velocity estimate evaluated at a $200 \times 200$ grid where $x \in (-2.5, 2.5)$ and $y \in (-4, 4)$. The velocity estimate from the flow map can be computed by $\frac{dx}{dt} = \frac{d}{dt}\Phi_0(\vec{x})$, $\vec{x} = [x, y]$.

#### 5.5 Controlling mapping smoothness

Another important aspect of flow maps parameterized by neural network is to control its "smoothness", since a small change in time should not result in a dramatic change in state variables. This can be quantified by computing the *Lipschitz constant*, $c$, defined as:

$$||\Phi_{t_0}(x) - \Phi_{t_1}(x)||_p \leq c||t_0 - t_1||_p. \tag{17}$$

Intuitively, the Lipschitz constant $c$ quantifies how fast output changes with respect to input under a specific choice of $p$-norm. For a fully connected layer, $f(x)$, defined by a weight matrix, $W$, and bias, $b$:

$$f(x) = Wx + b, \tag{18}$$

its Lipschitz constant can be computed directly by equation 17:

$$\begin{aligned} ||(Wx_1 + b) - (Wx_2 + b)||_p &\leq c||x_1 - x_2||_p \\ ||W(x_1 - x_2)||_p &\leq c||x_1 - x_2||_p \\ \frac{||W(x_1 - x_2)||_p}{||x_1 - x_2||_p} &\leq c, \end{aligned} \tag{19}$$

which end up being the operator norm of $W$, whose exact value will depend on the choice of $p$-norm. When $p = 1$ and $p = \infty$, the Lipschitz constant would be maximum column sum and row sum of $W$ respectively; and when $p = 2$, it would be the largest singular value of $W$.

For simplicity, we regularized the network by controlling its Lipschitz constant under the $\infty$-norm for its simplicity. With the choice, we can control the overall Lipschitz constant of the network by controlling the maximum row sum of each weight matrix since:

$$c(\Phi) \leq \prod_i c(W_i), \tag{20}$$

where $c(\Phi)$ denotes the Lipschitz constant of our network/flow map, and $W_i$ is the $i^{\text{th}}$ layer of the network [58, 59].

Motivated by the fact smoothness can be enforced by controlling row sums of weight matrices, we also improved network smoothness through weight decay, which penalizes large entries in weight matrices during optimization. Another route to increase network smoothness is to construct the network such that its derivative is continuous, which can be obtained most easily by using continuous activation functions such as the Gaussian error linear unit (GeLU) instead of the rectified linear unit (ReLU).

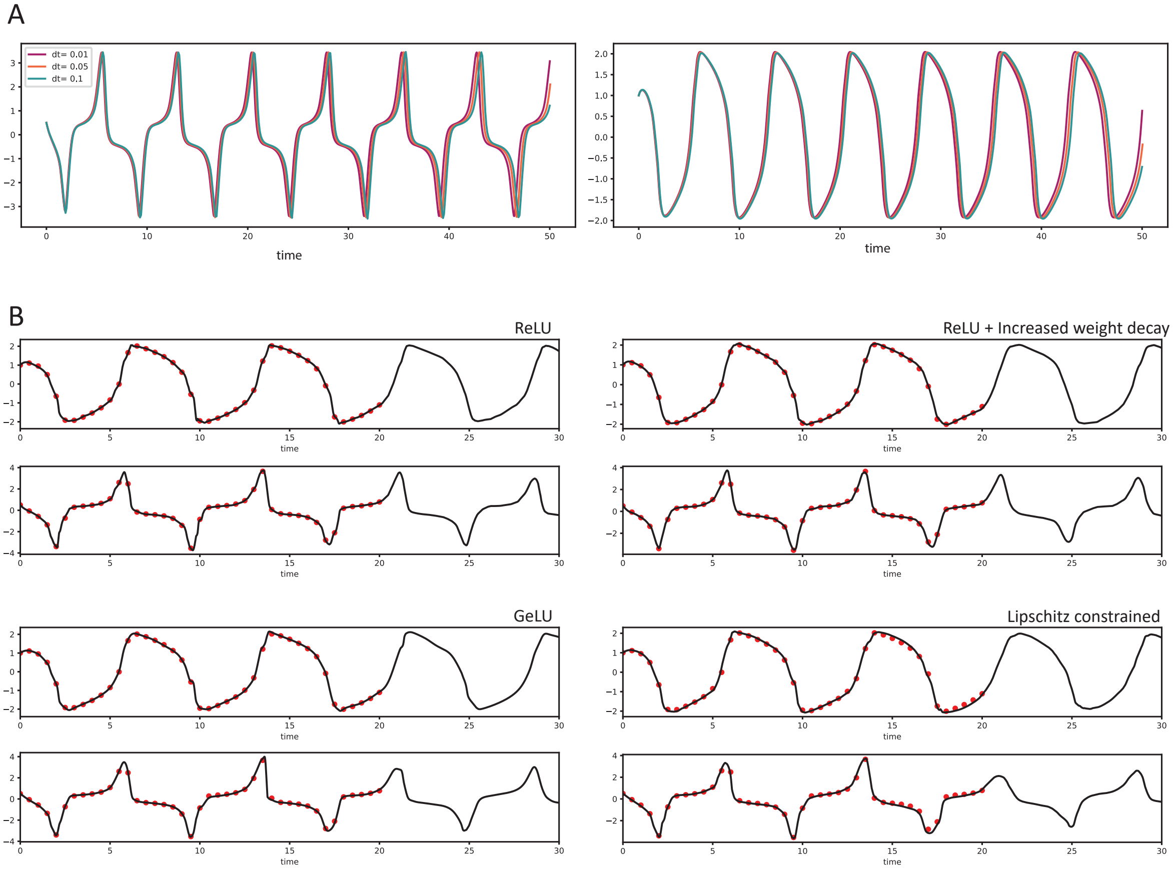


Figure S1: A: Time series generated with different $\Delta t$ using previously proposed semigroup loss. B: Time series generated with one forward pass with different regularization.

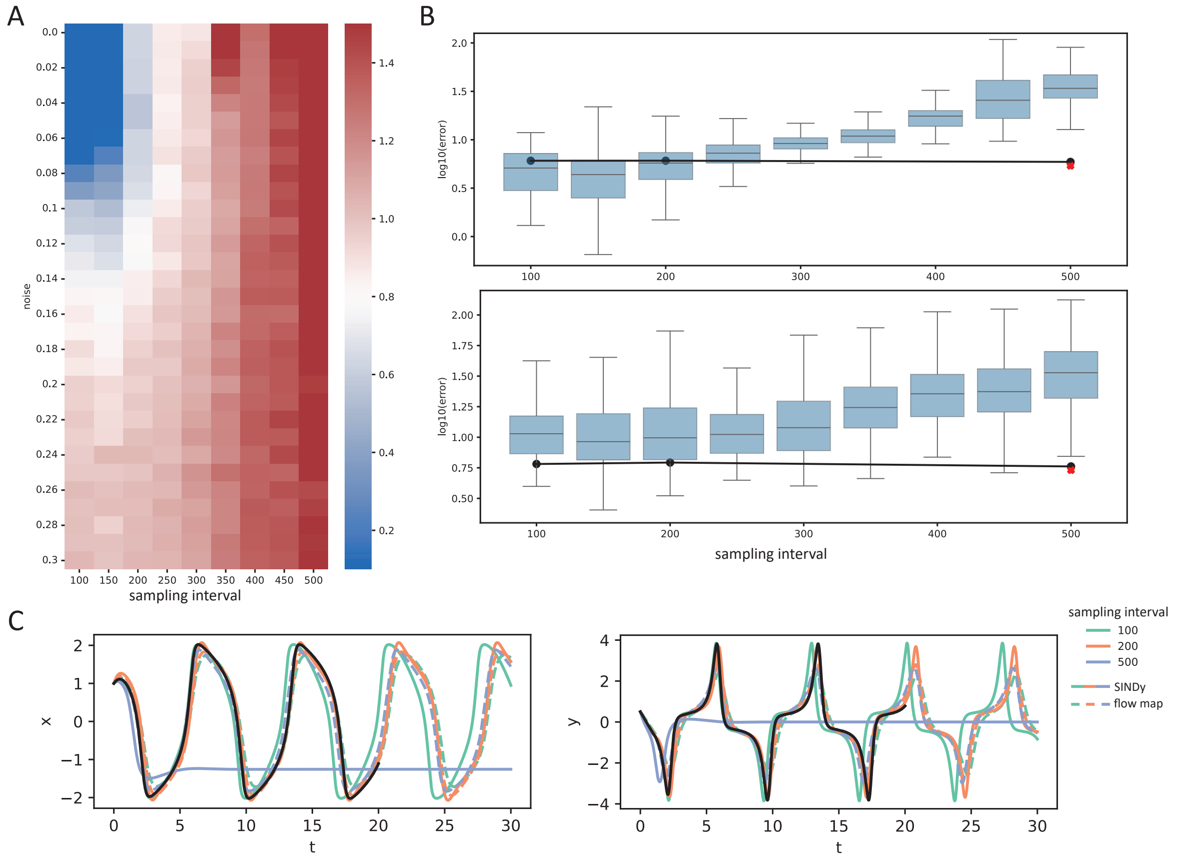


Figure S2: A: Error of the reconstructed Van der Pol oscillator via SINDy under different noise level and sampling interval. B: Boxplot showing the error of the reconstructed Van der Pol oscillator via SINDy under a noise level of 0.1 (top) and 0.3 (bottom). Line plot shows the error from the flow map reconstruction under the same condition. Red cross indicates the error from flow map trained with zero noise. C: Visualization of the reconstructed system via SINDy and flow map under different sampling interval with a noise level of 0.1.

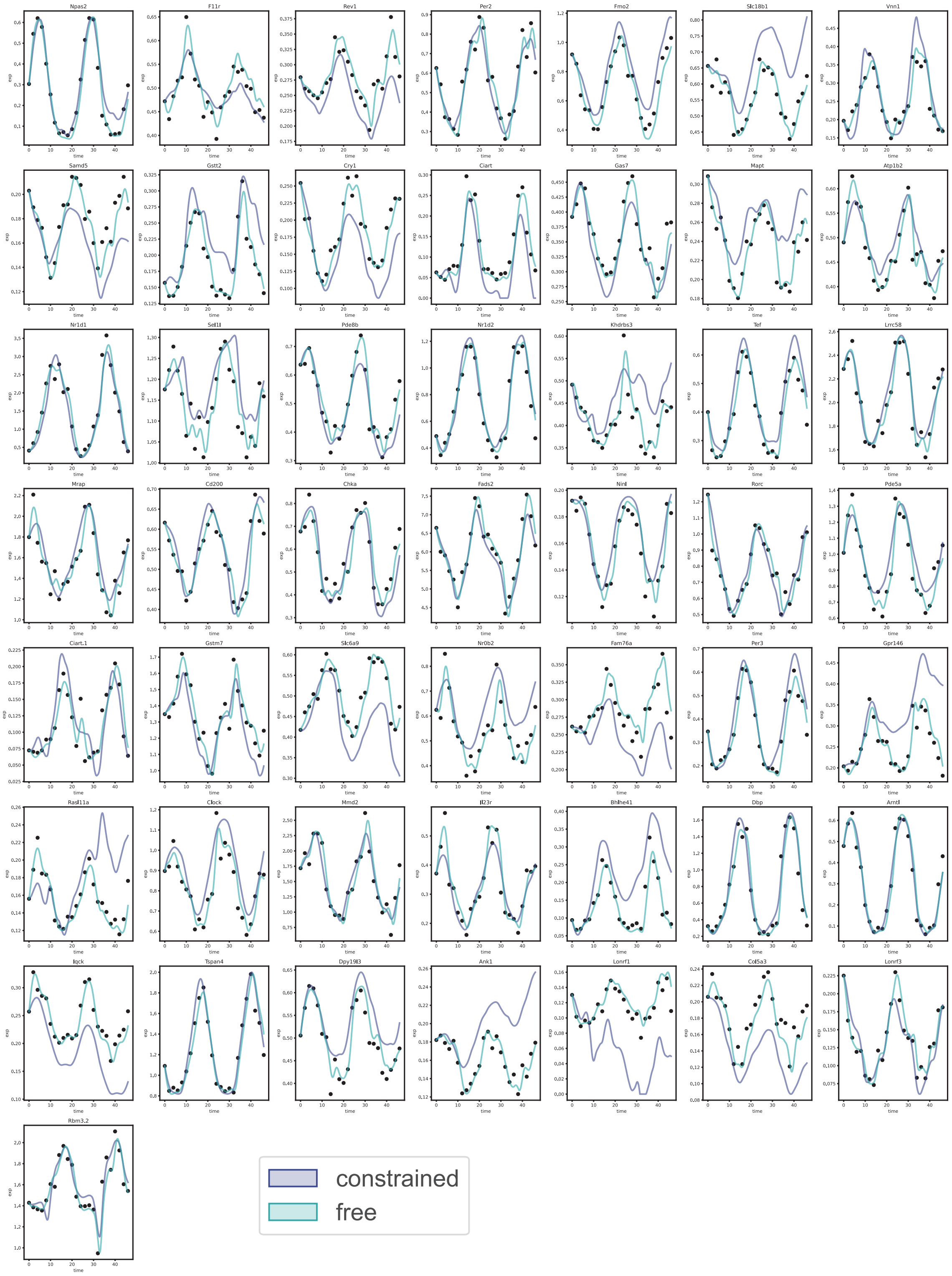


Figure S3: Reconstructed time series of all feature genes.

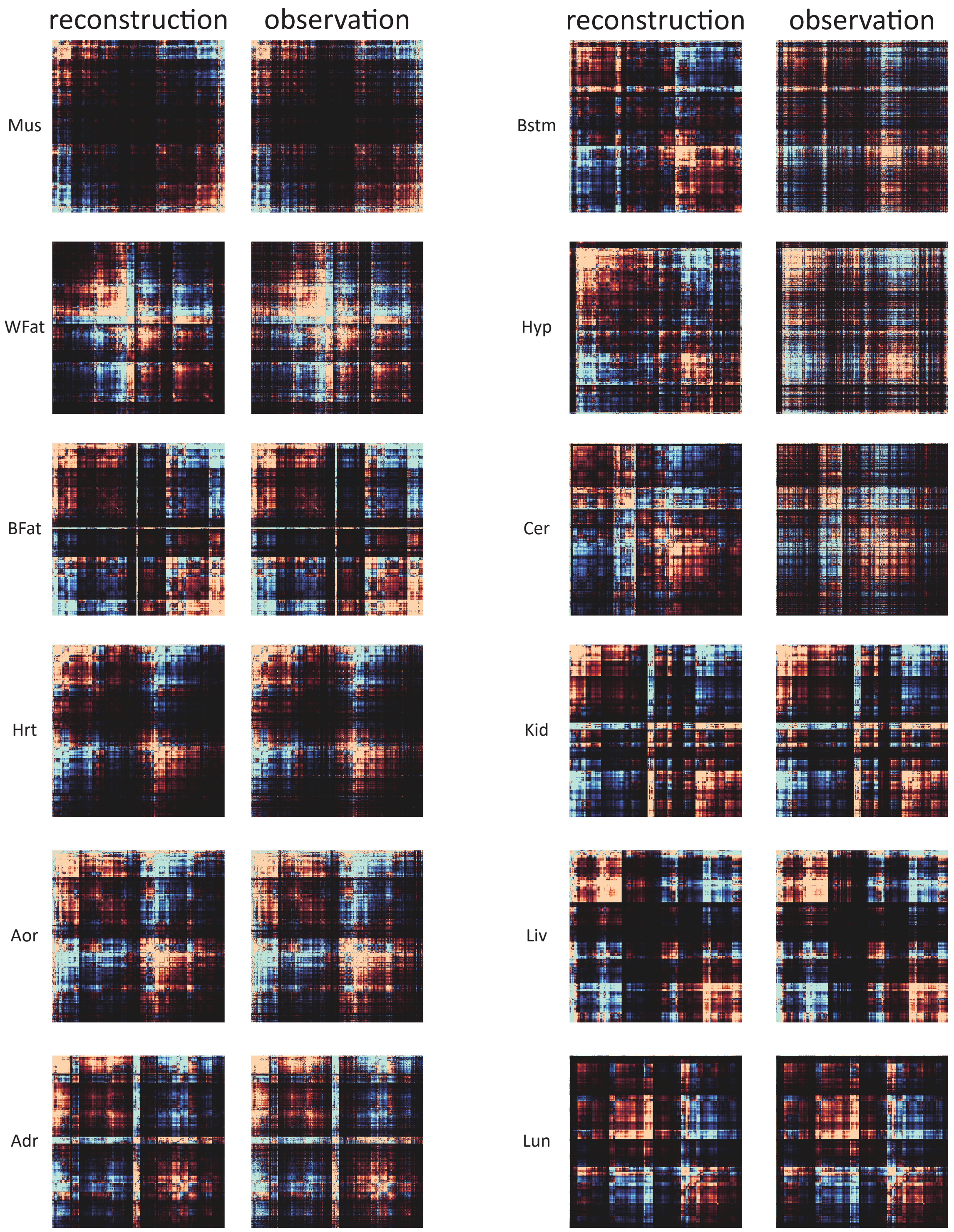


Figure S4: Observed and reconstructed covariance structure using the top 500 circadian genes across all organs.

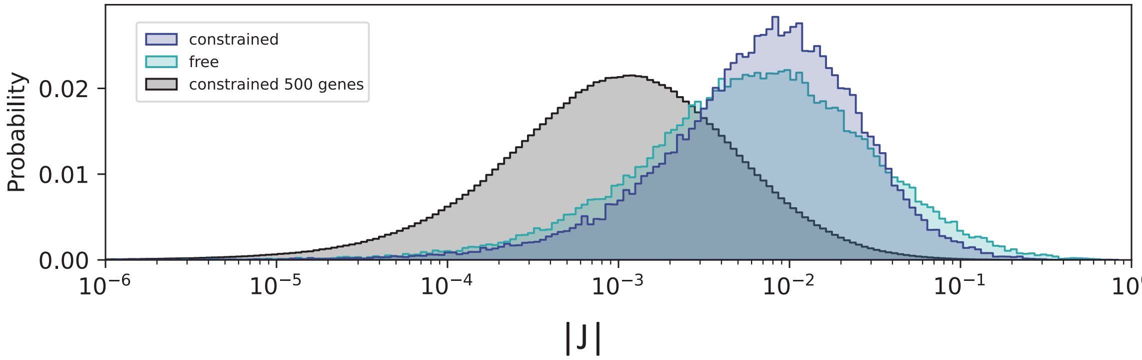


Figure S5: Probability distribution of the magnitude of the Jacobian matrix across all measured state in the adrenal gland.
